# Functional Connectivity Patterns Underlying the Experience of Auditory Verbal Hallucinations in Patients with Schizophrenia

**DOI:** 10.1101/062992

**Authors:** Parekh Pravesh, John P. John, Sangeetha Menon, Harsha N. Halahalli, Bindu M. Kutty

## Abstract

Schizophrenia is characterized by functional connectivity aberrations between brain regions that mediate different cognitive processes. The characteristic symptoms of schizophrenia such as delusions, hallucinations, passivity experiences etc. are suggested to reflect a disordered self-awareness. In the present study, we used a novel fMRI paradigm, the ‘Hallucination Attentional Modulation Task (HAMT)”, to examine the functional connectivity patterns underlying the experience of auditory verbal hallucinations in contrast to the patterns associated with processing of visual stimuli. We found that there was substantial overlap amongst healthy (*n*=8) and schizophrenia (*n*=6) subjects with respect to the functional connectivity patterns during the ‘free attention’ and ‘visual attention’ conditions of the paradigm. In patients with schizophrenia having continuous auditory verbal hallucinations, the connectivity between the bilateral superior parietal lobules and bilateral thalami were stronger during the ‘hallucination attention’ condition. These results provide preliminary leads that link auditory verbal hallucinations to an underlying disorder of self-agency.

## 1 INTRODUCTION

Schizophrenia is a complex psychiatric disorder characterized by symptoms such as delusions, hallucinations, passivity experiences etc. that are thought to arise from an aberrant self-awareness (Blakemore and Frith, 2003). Various symptom dimensions of schizophrenia have been linked to focal or generalized brain disconnection, such that it has come to be recognized as a disconnection syndrome (Friston and Frith, 1995). One of the most unique characteristic of schizophrenia, reported by around 75% of sufferers, is the experience of auditory verbal hallucinations (AVH) (Nayani and David, 1996). This is typically perceived as continuous conversations referring to the patient in third person or as a running commentary. Several theories have been put forth to explain the neurobiology of hallucinations including both “top-down” and “bottom-up” postulations (Feinberg, 1978; Frith and Done, 1988; Manford and Andermann, 1998; West, 1962). “Top-down” formulations such as the one by (Feinberg, 1978) propose that thoughts are perceived as hallucinations due to confusion regarding its source, thereby experiencing it as ‘alien’. On the other hand, “bottom-up” postulations highlight the role of impairment in sensory modalities in the causation of hallucinations (West, 1962). [See (Aleman and Vercammen, 2013) for a discussion on the top-down and bottom-up formulations of hallucinations].

There is a substantial body of functional neuroimaging studies that have examined the neural correlates of auditory hallucinations in schizophrenia. In a landmark Positron Emission Tomography (PET) study published in *Nature* carried out on 5 subjects with schizophrenia, Silbersweig et al., (1995) reported activations in subcortical nuclei, limbic structures, paralimbic regions as well as the sensory association cortices during the experience of auditory hallucinations. Since then, there have been many functional neuroimaging studies that have attempted to probe the neurobiology of AVH in schizophrenia. These include studies carried out using EEG (e.g., (Jardri et al., 2009)), PET (e.g., (McGuire et al., 1996, 1995)), structural MRI (e.g., (Gaser et al., 2004)), functional MRI (fMRI) (Allen et al., 2007; Amad et al., 2014; Dierks et al., 1999; Fu et al., 2001; Hoffman et al., 2008; Lawrie et al., 2002; Rolland et al., 2014; Shergill et al., 2001, 2001, 2000; Sommer et al., 2008; Vercammen et al., 2010), and DTI (e.g., (Hubl et al., 2004)). There is wide variability between these studies with respect to *(a) sample sizes* [from case reports like (Shergill et al., 2001), to sample sizes ranging from 4 (Lennox et al., 2000) to up to 27 subjects (Shinn et al., 2013)]; *(b) use of control group* [without control group (Dierks et al., 1999; Hoffman et al., 2008); healthy subjects as control (Lawrie et al., 2002); schizophrenia subjects without AVH as control (McGuire et al., 1995)]; *(c) design of task* [resting state (Vercammen et al., 2010), button-press to indicate onset of hallucination (Jardri et al., 2009), acoustical stimulation (Dierks et al., 1999) etc.]; and *(d) analyses methods* (between-group comparisons of whole-brain or regional activations (Fu et al., 2001; Sommer et al., 2008); functional connectivity analysis (Amad et al., 2014; Vercammen et al., 2010) etc.]. [See (Allen et al., 2008) for a comprehensive list of studies that have probed the neurobiology of hallucinations in schizophrenia].

Given such methodological heterogeneity across the studies, it is not surprising that there are substantial differences in the results that have emerged so far with respect to the brain regions/networks underlying hallucinations. Various brain regions have been linked by these studies to auditory hallucinations in schizophrenia: e.g., Heschl’s gyrus (primary auditory area); Broca’s area (motor speech area); bilateral hippocampus/ parahippocampal gyrus; ventral striatum; lenticular nuclei, anterior cingulate; orbitofrontal cortex; posterior cerebellar cortex, thalamus, middle temporal cortex, nucleus accumbens; right homologue of Broca’s area, bilateral insula; temporo-parietal junction etc. (Allen et al., 2007; Dierks et al., 1999; Hoffman et al., 2007; Shergill et al., 2000; Silbersweig et al., 1995; Sommer et al., 2008). Heschl’s gyrus has been one of the most commonly implicated brain regions; however there are many studies as well, that have not reported an association between activation in this brain region and auditory hallucination (e.g., (Silbersweig et al., 1995; Sommer et al., 2008)) In a recent meta-analysis, (Jardri et al., 2011) reported increased activation likelihoods in the bilateral neural networks involving Broca’s area, anterior insula, precentral gyrus, frontal operculum, middle and superior temporal gyri, inferior parietal lobule, and hippocampus/parahippocampal region in patients with schizophrenia having auditory hallucinations.

The above studies have attempted to capture the patterns of activations that are associated with AVH. However, given the premise that schizophrenia is a disorder of functional connectivity (Friston, 1999, 1998; Friston and Frith, 1995), it is of importance to look at the patterns of connectivity that underlie hallucinations. While most of the activation studies have adopted a functional segregation approach (the idea that different parts of the brain perform different functions), functional connectivity studies are based on a functional integration perspective (the idea that different parts of the brain function together to perform a particular task). Since complex cognitive and perceptual processes that are inherent to normal conscious awareness are the result of distributed processing involving connections between various brain regions, the examination of functional connectivity (FC) aberrations between different brain regions would provide a better understanding of network dysfunction underlying aberrant conscious experiences such as AVH.

The few fMRI studies that have attempted to examine connectivity disturbances associated with AVH have used resting state functional connectivity (rsFC) analysis (Amad et al., 2014; Gavrilescu et al., 2010; Rolland et al., 2014; Shinn et al., 2013; Vercammen et al., 2010). These studies have generated varied observations such as reduced interhemispheric connectivity in both primary and secondary auditory cortices in patients with hallucinations (Gavrilescu et al., 2010); reduced FC between the left temporo-parietal junction and the right homotope of the Broca’s area in patients with schizophrenia (Vercammen et al., 2010); increased FC between Heschl’s gyrus and left fronto-parietal regions and decreased FC between Heschl’s gyrus and right hippocampal formation as well as mediodorsal thalamus in patients with lifetime history of auditory hallucination versus those who did not (Shinn et al., 2013); increased seed based FC between the hippocampal complex and thalamus in patients with auditory hallucinations (Amad et al., 2014); increased FC of nucleus accumbens with left superior temporal gyrus, cingulate gyri, and the ventral tegmental area in patients with auditory hallucinations (Rolland et al., 2014); and lower FC between left temporal cortex and left dorsolateral prefrontal cortex in patients with schizophrenia (Lawrie et al., 2002). Vercammen et al., (2010) and Lawrie et al., (2002) reported an inverse relationship between the rsFC, predominantly in the frontal and temporal regions, and severity of auditory hallucinations, while Shinn et al., (2013) reported a positive relationship between rsFC and hallucination severity. Raij et al., (2009) have reported a positive correlation between subjective reality of the AVH and hallucination-related activation strength of the inferior frontal gyri (IFG) as well as the coupling between IFG, ventral striatum, auditory cortex, right posterior temporal lobe and the cingulate cortex. This study has examined functional connectivity using a single seed (IFG)-based approach using Psychophysiological Interactions (PPI) analysis in Statistical Parametric Mapping (SPM)-2 at a liberal statistical threshold uncorrected for multiple comparisons. Thus, given the methodological differences across these studies, no definitive conclusions regarding network dysfunction associated with AVH can be derived from these results, beyond inferring that there may be an impairment of fronto-temporal connectivity, especially in the left hemisphere.

A majority of the previous studies have not attempted to carefully choose patients with *continuous* auditory hallucinations, nor have they attempted to objectively verify whether these patients were experiencing hallucinations during the fMRI acquisition. Moreover, to the best of our knowledge, no study till date has attempted to examine the aberrations of functional connectivity between the various brain regions that are implicated in AVH in patients with schizophrenia during the experience of auditory hallucinations. In the ‘resting’ state, during fMRI acquisition, many patients may be experiencing auditory hallucinations while many others may not. Therefore, one cannot clearly infer from the results of resting state functional connectivity studies whether they reflect ‘trait’ abnormalities of schizophrenia as opposed to ‘state’ abnormalities underlying the experience of AVH. Task designs that involve button press to indicate onset of hallucination (e.g., (Raij et al., 2009; Silbersweig et al., 1995)) would result in unequal lengths of the time series which can influence the results of connectivity studies. Therefore, there is a need to develop approaches that are designed to examine the brain network dysfunction underlying the experience of AVH using appropriately designed paradigms in selected individuals with schizophrenia having continuous AVH.

In this paper, we present the results of a pilot study using a novel fMRI paradigm which we refer to as “Hallucination Attentional Modulation Task (HAMT)” during which monochromatic visual patterns changing at random intervals were presented on the screen. The task was comprised of five blocks, each containing three conditions, wherein the subject was instructed to focus attention on (a) nothing in particular (free attention condition), (b) the changing visual stimuli (visual attention condition) and (c) hallucinations (schizophrenia subjects) (hallucination attention condition) respectively. Such a paradigm permits the study of the effect of attentional modulation on functional connectivity in patients with schizophrenia having continuous auditory hallucinations by drawing their attention towards and away from the hallucinations. The paradigm also permits the study of processing of a true percept, i.e., visual stimuli in healthy subjects, and the effect, if any, of continuous auditory hallucinations on this in patients with schizophrenia. Most importantly, by including a subjective rating of the experience of AVH after every block of the hallucination attention condition (see Methods section), the paradigm provides an objective method of selecting blocks which were associated with the experience of hallucination, for further analyses. We performed this pilot study in a small sample of patients with recent-onset schizophrenia who were experiencing continuous AVH, as well as in healthy subjects. We carried out FC analysis between *a priori*-specified regions of interest (ROIs) (see Table 1) in patients with schizophrenia having continuous AVH during the performance of the HAMT. Given the premise that the experience of auditory hallucinations is possibly due to a deficit in differentiating between self and non-self (Blakemore et al., 2002; Blakemore and Frith, 2003; Feinberg, 1978; Feinberg and Guazzelli, 1999; Frith, 1987; Frith and Done, 1988), we hypothesised that patients experiencing AVH would show aberrant connectivity between brain regions involved in self-agency apart from those involved in auditory processing. We also hypothesized that brain regions that are involved in normal conscious awareness of visual stimuli would show significant connections, which may be altered in patients with continuous AVH.

## 2 MATERIALS AND METHODS

The study was carried out at the National Institute of Mental Health and Neurosciences (NIMHANS), Bangalore, India with due approval from the Institute Ethics Committee, thus conforming to the ethical standards laid down in the 1964 Declaration of Helsinki. Written informed consent was obtained from all participants (and their legally qualified representatives in case of patients with schizophrenia) before enrolling them for the study.

### 2.1 Study Samples

The study samples for this exploratory pilot study consisted of 6 patients with schizophrenia (5 males) having continuous AVH, recruited from the out-patient department of NIMHANS by purposive sampling and 8 healthy subjects (4 males) (recruited from staff and students of NIMHANS). The diagnosis of schizophrenia was made through consensus between a consultant psychiatrist who used DSM-IV-TR criteria (American Psychiatric Association, 2000) and a trained research assistant who performed a screening using Mini-International Neuropsychiatric Interview (MINI) Plus (Sheehan et al., 1998). All subjects were right handed (as determined by Edinburgh handedness inventory (Oldfield, 1971)), between the age of 17-50 years, and had a Hindi mental state examination (HMSE) (Ganguli et al., 1995) score of ≥ 23 (in order to ensure that all subjects had an adequate level of cognitive functioning). All subjects were native speakers of two of the South Indian Dravidian languages (Kannada or Tamil) and were comfortable in reading their preferred language (or English). The patients with schizophrenia who were recruited for the study, had continuous auditory hallucinations defined using modified criteria from PSYRATS (Haddock et al., 1999) on the basis of presence of continuous auditory hallucinations for more than 50% of the awake time for at least the previous 2 weeks. Healthy subjects were also screened using MINI Plus to rule out any past or present psychiatric disorders. History of psychiatric disorders in first degree relatives, medical or neurological conditions requiring regular medications, and current exposure to psychotropic medications were ruled out by an unstructured interview before recruitment into the study.

The schizophrenia sample had a mean age of 33.67 years (s.d. = 7.99; range: 22-40) and a mean of 11.33 years of education (s.d. = 3.93; range: 5-16). The mean WAIS-III score was 12 (s.d. = 5.01), and the mean HMSE score was 30.17 (s.d. = 0.75). Five of the patients were diagnosed as having paranoid schizophrenia while one patient was diagnosed as having schizoaffective disorder, depressed subtype. The mean age of onset of schizophrenia was 29.16 years (s.d. = 7.67; range: 20-39) and the mean duration of illness was 54.66 months (s.d. = 54.31; range: 6-120 months). At the time of recruitment into the study, two patients were neuroleptic naïve, one was neuroleptic free, and the other three were on regular medications. The mean scores on the Scale for the Assessment of Positive Symptoms (SAPS) and Scale for the Assessment of Negative Symptoms (SANS) were 51.82 (s.d. = 22.47) and 23 (s.d. = 14.65) respectively. On the PSYRATS scale, the mean score was 34.33 (4.54 s.d.) and 15.5 (8.38 s.d.) for hallucinations and delusions respectively. None of the patients were having significant drug-related side effects or movement disorder at the time of their participation in the study. The healthy sample had a mean age of 27 years (s.d. = 4.20; range: 20-34) and a mean of 13.87 years of education (s.d. = 4.12; range: 9-20 years). The mean WAIS-III rating was 13 (s.d. = 3.07) and the mean HMSE score was 30.87 (s.d. = 0.35).

### 2.2 Experimental design

The fMRI experiment consisted of three conditions implemented in a block design pattern using E-Prime stimulus presentation software, version 1.0 (PST Inc, Philadelphia, USA, www.pstnet.com/) operating within the Integrated Functional Imaging System (IFIS-SA) (Invivo, Orlando, Florida, USA, www.invivocorp.com) running on Windows XP. The three conditions involved in the Hallucination Attentional Modulation Task (HAMT) were: ‘Free Attention (FA)’, ‘Visual Attention (VA)’, and ‘Hallucination Attention (HA)’.

The task involved the presentation of a series of abstract monochromatic visual images that changed at unpredictable intervals. Images were 200×200 pixels in dimension and were presented at a resolution of 72 pixels per inch. These images were created using GNU Image Manipulation Program (GIMP) 2.4.7, using the fill pattern tool, creating six distinct patterns which were clearly distinguishable from each other. The task consisted of five blocks, with each block having three conditions (two in case of healthy subjects). Each condition began with an instruction slide displayed for 9 seconds and ended with an end-of-condition response collection slide. Each condition lasted (including task instruction and response collection) 42 seconds (Figure 1).

During the “free attention (FA)” condition, the subjects were instructed to keep their eyes fixed on the centre of the screen and to not pay attention to anything in particular. Changing abstract patterns were presented on the screen for a duration ranging from 2 to 18 seconds each (in a predetermined pseudorandom order). The number of images displayed varied from 4-9 depending on the stimulus duration, with the stimulus presentation lasting for 24 seconds. At the end of the condition, a response collection slide was presented for 9 seconds, asking the subjects to press any button of their choice. In the “visual attention (VA)” condition, subjects were asked to keep their eyes fixed on the centre of the screen and to count the number of times the picture on the screen changed. Attentional engagement was assessed at the end of the condition by presenting the subjects with a response collection slide asking them the number of times the images changed. Subjects were instructed to indicate their choice by button press from among the three choices presented. In case of patients with schizophrenia, an additional condition was introduced. During this “hallucination attention (HA)” condition, patients were asked to keep their eyes fixed on the centre of the screen but to pay attention to their AVH (if any). At the end of the condition, a slide with three choices were presented to the subjects: “No voices were heard - Press index finger; Voices heard, but for less than 50% of time - Press middle finger; Voices heard for more than 50% of time - Press ring finger”, and the appropriate response was recorded. 4 of the 6 schizophrenia subjects reported ‘hearing voices’ for more than 50% of the time during the hallucination attention condition of all the 5 blocks (classified as AVH+ subjects), while the remaining 2 reported not experiencing hallucinations during ≥ 2 of the 5 blocks.

**Figure 1:**
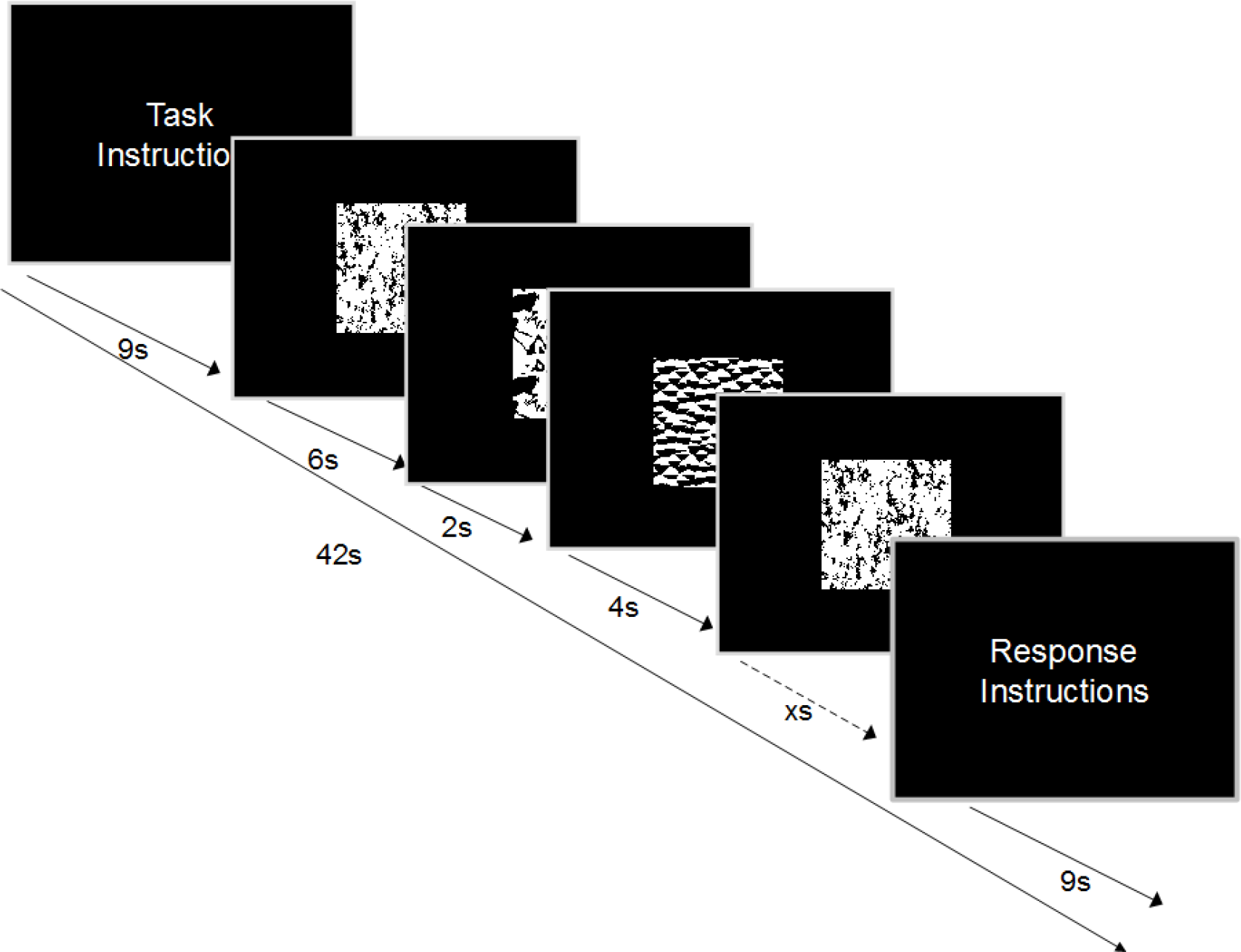
Hallucination Attentional Modulation Task (HAMT): fMRI experiment design. The task involved presentation of abstract monochromatic images that changed at unpredictable interval. During the free attention (FA) condition, subjects were asked to keep their eyes fixed on the centre of the screen and not pay attention to anything in particular. During the visual attention (VA) condition, subjects were asked to count the number of times the image changed on the screen. During the hallucination attention (HA) condition (in case of patients with schizophrenia), subjects were asked to pay attention to their auditory verbal hallucinations, if any. Each block was preceded by instructions and ended with end-of-block response collection (see text for details).

### *2.3* Image Acquisition

Magnetic resonance imaging (MRI) was performed on a Philips Achieva 3.0 T scanner using an 8-channel phased array head coil. Head movements were minimized by applying a band over the forehead during the scanning procedure. Single-shot echo-planar images (EPI) were acquired at an in-plane resolution of 2×2×6 mm with axial, sequential, contiguous 6 mm slices using a SENSE 8 acceleration; TR/TE, 3,000 ms/35 ms; flip angle, 90°; and EPI factor of 59. A volume shim covering parts of the frontal and temporal lobes was used to minimize distortions induced by air-tissue interfaces adjacent to the sinuses. The acquisition was preceded by two dummy volumes that were discarded. A high-resolution (1×1×1 mm voxel size) T1-weighted MRI volume data set of the whole brain was also obtained using the same head coil for each subject [Magnetization Prepared Rapid Gradient Echo (MP-RAGE sequence) with TR 8.2 ms, TE 3.8 ms, flip angle 8°, and sense factor 3.5]. All images were stored in DICOM format.

### *2.4* ROIs for functional connectivity analysis

ROIs for FC analysis were defined *a priori* based on previous reports of brain regions that have been linked to AVH (Vercammen et al., 2010); (Rolland et al., 2014); (Shinn et al., 2013). Additionally, primary and secondary visual areas of the brain (BA17, BA18, and BA19) were also included, given the nature of the task. The ROI set comprised of 16 bilateral regions (Table 1), defined using the human brain atlas in WFU PickAtlas (Lancaster et al., 2000, 1997; Maldjian et al., 2003) toolbox in MATLAB (MathWorks Inc, Natick, Maryland, USA, www.mathworks.com).

**Table 1:** List of ROIs included for functional connectivity analysis including their four letter abbreviation and their centroid location. ROIs were defined using the Human Brain Atlas in WFU PickAtlas toolbox in MATLAB.

| Region of Interest | Label | L/R | Centroid |  |  |
| --- | --- | --- | --- | --- | --- |
|  |  |  | X | Y | Z |
| Amygdala | Amyg | L | -22.93 | -4.03 | -18.78 |
|  |  | R | 23.73 | -3.83 | -18.73 |
| Anterior Cingulate | AnCG | L | -7.74 | 32.7 | 9.09 |
|  |  | R | 8.44 | 32.46 | 8.62 |
| Broadman Area 17 | BA17 | L | -11.38 | -92.01 | -3.72 |
|  |  | R | 12.4 | -92.18 | -4.19 |
| Broadman Area 18 | BA18 | L | -18.42 | -88.19 | -1.32 |
|  |  | R | 19.06 | -88.23 | -1.28 |
| Broadman Area 19 | BA19 | L | -31.51 | -79.41 | 11.57 |
|  |  | R | 32.97 | -79.54 | 11.4 |
| Inferior Frontal Gyrus | InFG | L | -44.22 | 24.08 | 3.91 |
|  |  | R | 45.55 | 24.14 | 4.07 |
| Inferior Parietal Lobule | InPL | L | -47.63 | -43.88 | 39.56 |
|  |  | R | 48.99 | -44.11 | 39.65 |
| Insula | InSL | L | -38.61 | -7.43 | 9.46 |
|  |  | R | 39.76 | -7.61 | 9.42 |
| Medial Frontal Gyrus | MeFG | L | -8.36 | 28.67 | 26.64 |
|  |  | R | 9.15 | 26.03 | 29.09 |
| Middle Frontal Gyrus | MiFG | L | -36.84 | 28.4 | 29.05 |
|  |  | R | 38.27 | 28.34 | 29.28 |
| Nucleus Accumbens | NuAc | L | -13.04 | 7.24 | -11.96 |
|  |  | R | 12.36 | 9.06 | -11.16 |
| Precuneus | PCun | L | -14.23 | -64.96 | 40.71 |
|  |  | R | 14.9 | -64.88 | 40.62 |
| Parahippocampal Gyrus | PaHG | L | -24.95 | -24.83 | -16.08 |
|  |  | R | 25.86 | -24.84 | -16.18 |
| Superior Parietal Lobule | SuPL | L | -25.31 | -63.58 | 54.33 |
|  |  | R | 26.77 | -63.42 | 54.37 |
| Superior Temporal Gyrus | SuTG | L | -50.33 | -17.06 | -1.98 |
|  |  | R | 51.28 | -16.92 | -2.05 |
| Thalamus | Thal | L | -12.39 | -19.5 | 7.57 |
|  |  | R | 12.7 | -19.48 | 7.62 |

### *2.5* Functional connectivity analysis

Image pre-processing and functional connectivity analyses were carried out using the Conn Functional Connectivity Toolbox 15.0h (Whitfield-Gabrieli and Nieto-Castanon, 2012) running in MATLAB environment with SPM 12b in the background. Conn implements an anatomical component-based noise correction method (aCompCor) (Behzadi et al., 2007) which has been previously shown to be useful for functional connectivity analysis by increasing the validity, sensitivity, and specificity of the analysis (Chai et al., 2012). Preprocessing of the images was carried out using the default pipeline consisting of realignment and unwarp, setting the origin of functional scans to (0,0,0), slice timing correction (interleaved bottom-up), setting the origin of structural scans to (0,0,0), structural segmentation and normalization (to MNI), functional normalization (to MNI), detection of outliers (using the artefact rejection toolbox method built-in) and finally functional smoothing. A Gaussian kernel of 9 mm full-width at half maximum (FWHM) was specified for functional smoothing.

Scrubbing parameters were estimated by setting the global signal z value threshold to 9 and subject motion threshold to 2mm. Global signal was not regressed and a band-pass of 0.008 to Inf was implemented during the denoising step. Confounds of white matter, CSF, realignment, scrubbing, and the main effects of each condition were included for linear regression in the denoising step. Percentage signal change (PSC) was used as the unit of analysis with signal from each ROI scaled to their own ROI-specific average BOLD signal. Time series for FC analysis was extracted from un-smoothed (i.e. normalized) functional images. Functional connectivity between any pair of ROIs was defined as the bivariate correlation between their respective functional time series. The correlation coefficients were converted to Fisher’s *Z* value to improve the normality of the data.

We studied the functional connectivity patterns between the above ROIs in healthy subjects and in patients with schizophrenia for FA, VA and HA conditions. Further, we also explored functional connectivity in patients who experienced hallucinations (AVH+) during the HA condition separately. Given that this is a pilot study, the limited sizes of the schizophrenia and healthy samples do not permit valid between group (HS vs. SZ or AVH+ vs. AVH-) comparisons. Therefore, one sample *t*-tests were performed to test whether the mean functional connectivity strength between pairs of ROIs was significantly non-zero at *α*<0.05 (FDR corrected). This would permit examination of functional connectivity patterns during the FA and VA conditions in both schizophrenia and healthy subjects as well as during the experience of AVH in patients with schizophrenia.

## 3 RESULTS

### *3.1* Visualization of results

A ‘seed level FDR’ was used to jointly evaluate whether the connectivity between each seed ROI and all the other ROIs showed any significant effect of interest. This was done by selecting all 32 ROIs and ‘*p*-FDR (seed-level correction)’ in Conn. The resulting connectivity graphs show connection thicknesses which are proportional to their *T* values (these figures are presented in the supplementary section). It should be noted that for a pair of seed and target, it is possible that the connection between them would pass the FDR correction when one is the seed but not the other way round. This is because the FDR correction would depend on all the other connections that are also involved for each seed. *Z* value of all significant connections were exported from Conn and connectivity graphs (with edges proportional to *Z* value) were created in Gephi (Bastian et al., 2009). These were, in turn, superimposed on a template brain to generate connectivity graphs similar to the output of Conn. Also, the colour of the nodes was assigned based on the number of connections of each seed ROI (or *degree* as per graph theory terminology). The ROIs were grouped based on the functional networks that are linked to the processing of the visual stimuli and the experience of AVH. Based on the strength of connections, we present some of the significant connections that show patterns that differentiate between the different conditions and the groups (Table 2). [vide Supplementary Tables 1-6 for a comprehensive list of significant connections in the three conditions in the healthy, schizophrenia and AVH+ subjects]

### *3.2* Free attention condition

During the FA condition, healthy subjects showed widespread connectivity between the ROIs. The ROIs that showed high degree of connectivity include those involving regions that constitute the *internal awareness network* (medial prefrontal cortex, precuneus and inferior parietal lobule); *external awareness network/fronto-parietal network* (middle and inferior frontal gyri, inferior parietal lobule); *salience network* (insula and anterior cingulate); *visual network* (BA 17, 18, 19), *auditory cortex* (bilateral superior temporal gyrus) and *bilateral parahippocampal gyri, bilateral thalamus* as well as *posterior parietal regions (superior parietal lobule)*. Most of the ROIs showed high *degree* of connections, as represented in Figure 2.

From figures 2a and 2b, it is quite obvious that patients with schizophrenia have substantially lower extent of connectivity between the ROIs as compared to the healthy subjects. Moreover, the ROIs showed lower *degree* of connectivity as compared to healthy subjects.

Overall, the strongest connections between the ROIs in the FA condition were similar amongst both healthy and schizophrenia subjects, involving the above mentioned networks/brain regions (Table 2). Both the groups showed strong connections between the visual network ROIs, given the changing visual stimuli in the background during fMRI acquisition. Nevertheless, the degree of most of the ROIs including the visual network ROIs was lower in patients with schizophrenia. The prominent differences in connectivity noted in patients with schizophrenia included weaker connection between right anterior cingulate and right medial prefrontal cortex as well as reduced connectivity between left inferior and middle frontal gyri, and bilateral visual association area (BA18). Conspicuously, the connection between left and right superior parietal lobules *(Schizophrenia: Z=1.0336, range of Z values of all the significant ROI-ROI connections in the FA condition: 0.0866-1.0860; Healthy subjects: Z=0.7765; range of Z values of all the significant ROI-ROI connections in the FA condition: 0.1303-1.1860)* as well as left and right thalamus *(Schizophrenia: Z=0.9325; Healthy subjects: Z=0.7824)* was substantially higher in patients with schizophrenia as compared to healthy subjects.

**Figure 2:**
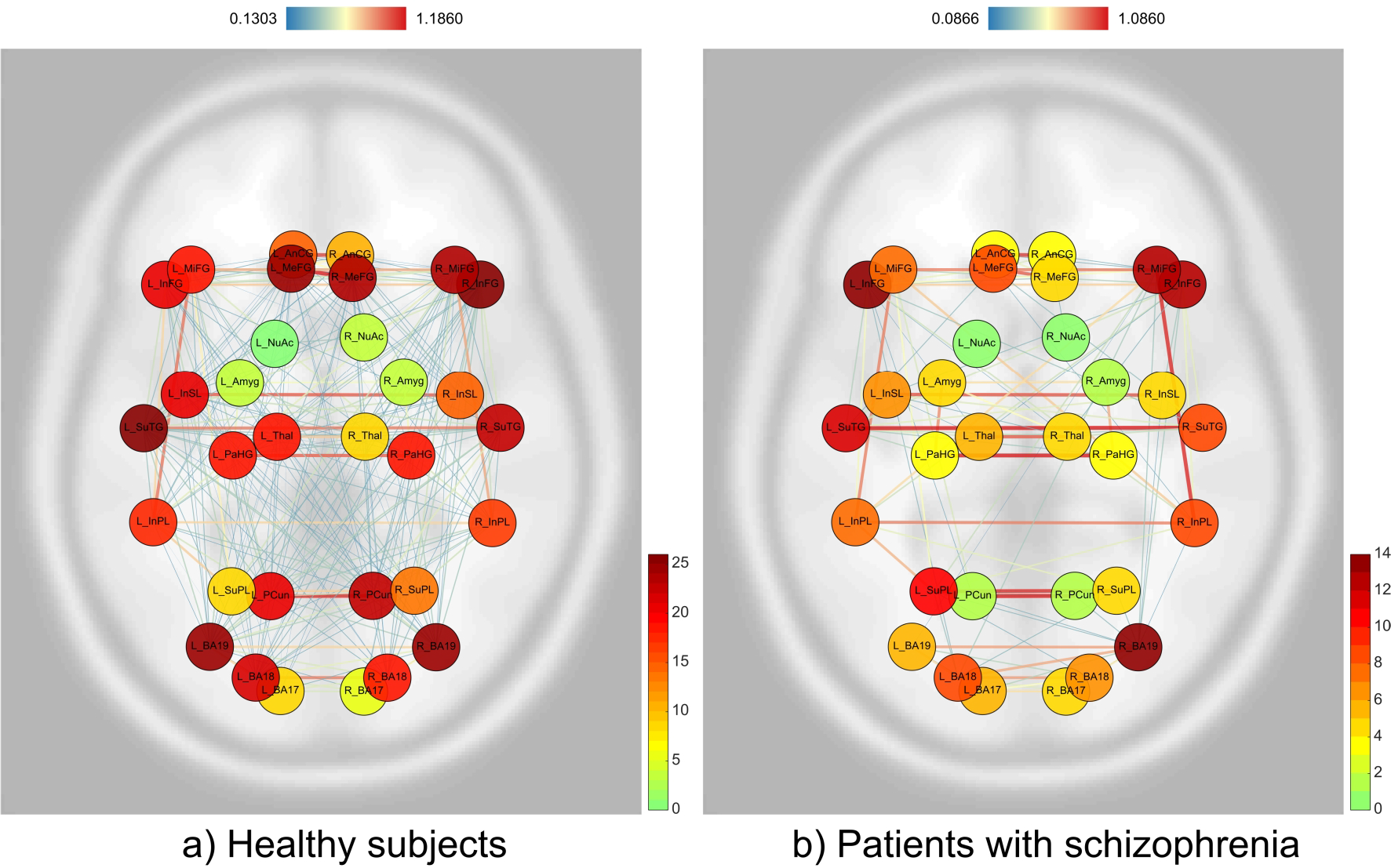
Connectivity graphs during free attention condition in a) healthy subjects; and b) patients with schizophrenia. The edge colours are proportional to *Z* values (colour bar on top) while the colour of the nodes corresponds to the number of connections (or degree) of each of the seed ROIs (colour bar on the right).

### 3.3 Visual attention condition

Even in the VA condition, the overall pattern of similarities and differences between healthy and schizophrenia subjects seen in the FA condition was noted (Figures 3a and 3b). The strongest connections between the ROIs were similar in both the groups. The visual network ROIs were also well-connected in both the groups. Nevertheless, the strength of connections and the degree of connectivity of the ROIs (including the ROIs in the visual network) were lower in the schizophrenia group, as was noted in the FA condition. Importantly, as was the case in the FA condition, the connection between left and right superior parietal lobules *(Schizophrenia: Z=1.1536, range of Z values of all the significant ROI-ROI connections in the VA condition: 0.1869-1.2676; Healthy subjects: Z=0.8093, range of Z values of all the significant ROI-ROI connections in the VA condition: 0.0991-1.2346)* as well as left and right thalamus *(Schizophrenia: Z=0.9923; Healthy subjects: Z=0.8263)* was substantially higher in patients with schizophrenia as compared to healthy subjects. Right anterior cingulate-medial prefrontal cortex, left BA18-BA19 connections, and bilateral BA18 connections were substantially weaker than that of the healthy subjects, similar to what was observed in the FA condition.

**Figure 3:**
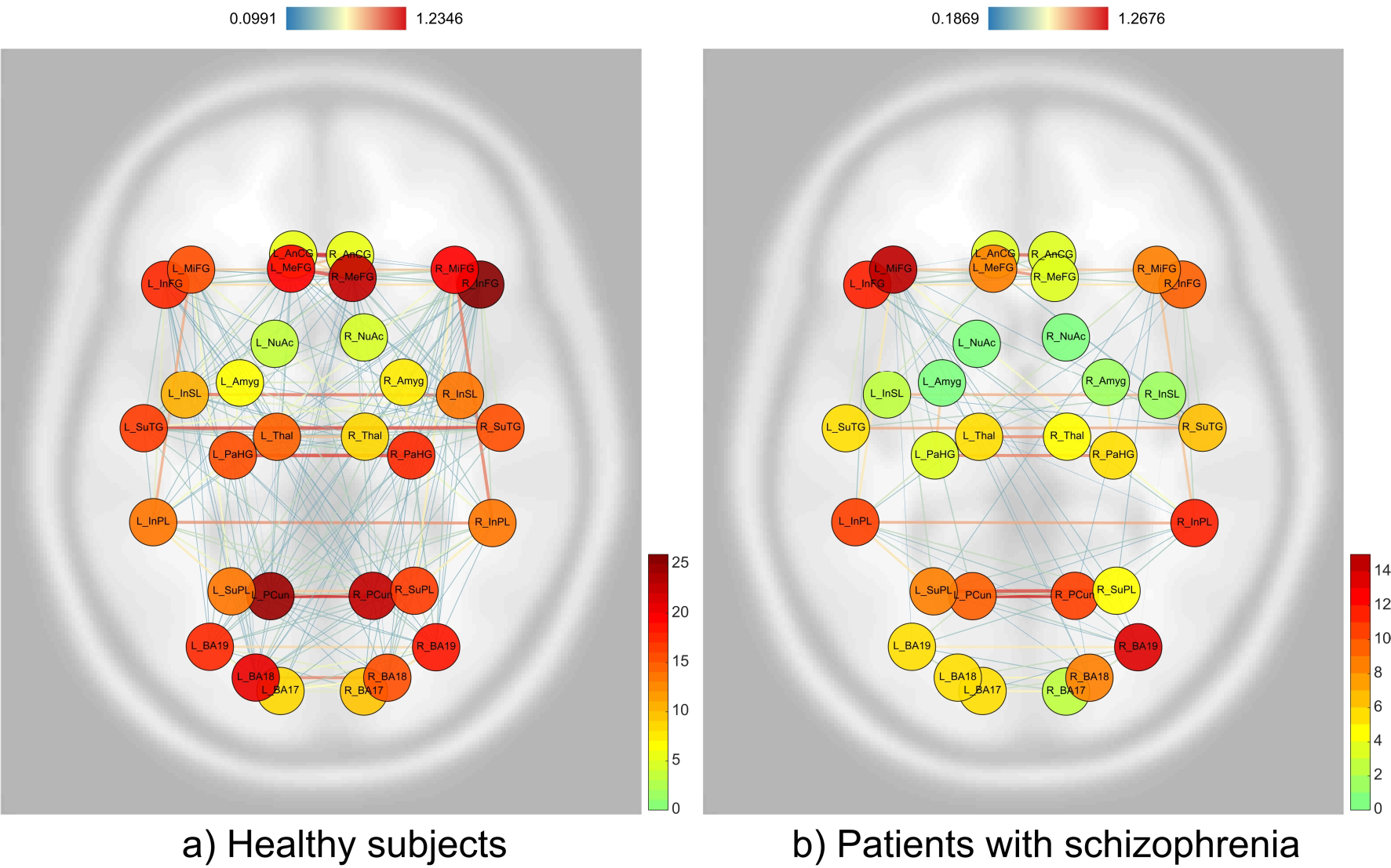
Connectivity graphs during visual attention condition in a) healthy subjects; and b) patients with schizophrenia. The edge colours are proportional to *Z* values (colour bar on top) while the colour of the nodes corresponds to the number of connections (or degree) of each of the seed ROIs (colour bar on the right).

### 3.4 Hallucination attention condition

As was the case with FA and VA in both healthy and schizophrenia subjects, high connection strengths were noted between bilateral medial prefrontal cortices, precuneus, anterior cingulate, insula, parahippocampal gyrus, superior temporal gyrus and the visual association cortices. Again, the most conspicuous difference that was noted between the hallucination attention condition in patients with schizophrenia in comparison to the FA condition in healthy subjects were the connections between left and right superior parietal lobules *(Schizophrenia: Z=1.0527, range of Z values: 0.1304-1.0828; Healthy subjects: Z=0.7765, range of Z values: 0.1303-1.1806)* as well as left and right thalamus *(Schizophrenia: Z=0.9190; Healthy subjects: Z=0.7824)*. The left superior parietal lobule was more strongly connected to the left inferior parietal lobule in patients with schizophrenia. As in the case of the FA and VA conditions, there was substantially weaker connection between the right anterior cingulate and medial prefrontal cortex; in addition, the bilateral inferior frontal gyri connection was also substantially reduced. The connections between the brain regions that constitute the fronto-parietal network were found to be substantially lower during the HA condition in comparison to the other conditions in both groups of subjects. The connection between the bilateral visual association cortices (BA18) was also noted to be lower, as was the case in the FA and VA conditions in patients with schizophrenia.

Functional connectivity analysis in the sub-group of 4 patients with schizophrenia who experienced AVH during the hallucination attention (HA) condition (AVH+) revealed an interesting pattern that underlined the findings of increased connections between left and right superior parietal lobules *(Z=1.0506)* and left and right thalamus *(Z=0.9993)* associated with AVH. As was noted with the overall schizophrenia sample, the left superior lobule was more strongly connected to the left inferior parietal lobule in the AVH+ sub-group. Uniquely in the AVH+ subjects, the bilateral precuneus connection became non-significant during the HA condition. The connections within the salience network were also noted to be either weaker or non-significant. As was the case with the overall sample of patients with schizophrenia, the fronto-parietal connections were substantially lower/non-significant in the AVH+ subjects; in addition, the connections within the visual network were also nonsignificant. Notably, the connection between the bilateral superior temporal gyri was comparatively weaker in the schizophrenia sample during the HA condition in comparison to the other conditions in both groups of subjects; in AVH+ subjects this connection was noted to be even weaker.

**Figure 4:**
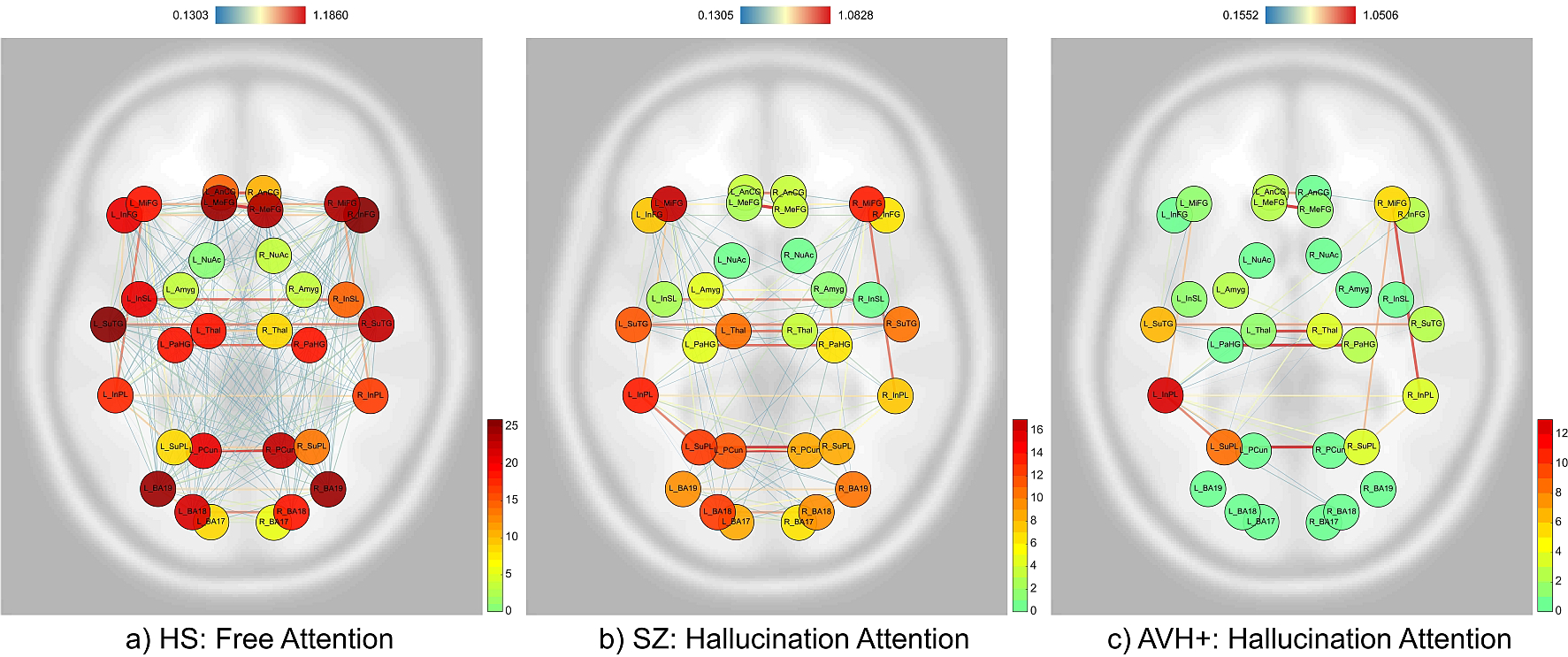
Connectivity graphs during a) free attention condition in healthy subjects; b) hallucination attention condition in patients with schizophrenia; and c) hallucination attention condition in AVH+ schizophrenia subgroup. The edge colours are proportional to *Z* values (colour bar on top) while the colour of the nodes corresponds to the number of connections (or degree) of each of the seed ROIs (colour bar on the right).

**Table 2:**
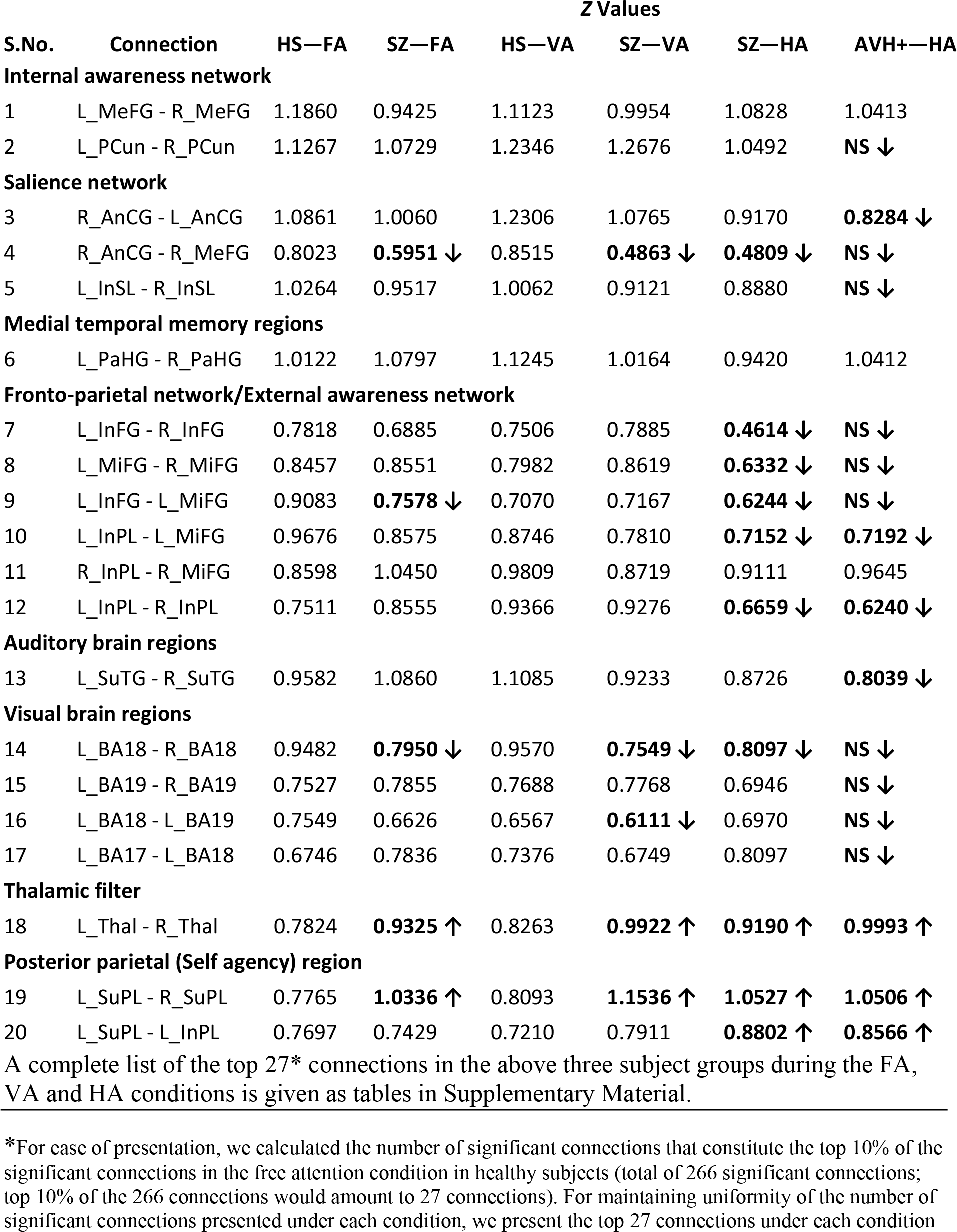
Selected brain regions that show patterns of significant connections that differentiate between conditions [free attention (FA), visual attention (VA) and hallucination attention (HA)] and subject groups [healthy (HS) schizophrenia (SZ) and patients who experienced hallucinations (AVH+)]. The brain regions are grouped under different functional networks/regions that are linked to processing of visual stimuli and the experience of auditory verbal hallucinations.

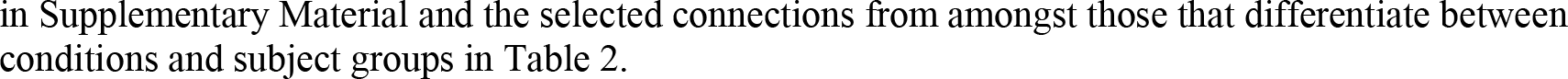

## 4 DISCUSSION

The objective of this pilot study was to examine the utility of a novel fMRI task which we have named “Hallucination Attention Modulation Task (HAMT)” in detecting functional connectivity aberrations associated with the experience of AVH. Certain internally consistent patterns have emerged from the study with respect to connections between brain regions during perception of a real visual stimulus in healthy and schizophrenia subjects as well as during perception of AVH in schizophrenia subjects. These observations highlight the potential utility of the above novel fMRI task for detecting functional connectivity aberrations underlying the experience of AVH.

The observation of a reduction in overall connections in the schizophrenia subjects (Figures 2,3, and 4) could provide support to the disconnection hypothesis of schizophrenia (Friston, 1999, 1998; Friston and Frith, 1995). Bilateral interhemispheric connections noted between various homologous brain regions during different conditions in both groups are in keeping with the basic organizing principles of brain functioning (see, for example, (Salvador et al., 2005)).

Since changing visual stimuli were presented across all the conditions, the brain regions that showed similar patterns of significant connectivity in both healthy and schizophrenia subjects during FA and VA conditions may be considered to be associated with the perception of real visual stimuli. These include, apart from the regions that constitute the *visual network* (BA 17, 18, 19), the *internal awareness network* (medial prefrontal cortex, precuneus) (Demertzi et al., 2013); *external awareness network/fronto-parietal network* (middle and inferior frontal gyri, inferior parietal lobule) (Demertzi et al., 2013); and *salience network* (insula and anterior cingulate) (Menon and Uddin, 2010). Additionally, brain regions that are involved with *memory* (parahippocampal gyri, amygdala) (Phelps, 2004; Takahashi et al., 2002), *self-agency* (superior parietal lobule) (MacDonald and Paus, 2003) and *sensory filtering* (thalamus) (McCormick and Bal, 1994) showed significant functional connectivity during FA and VA conditions in both healthy and schizophrenia subjects.

The brain regions that showed substantially stronger connections in patients with schizophrenia, especially in the AVH+ subjects, over and above those regions involved in perception of the real visual stimuli, may be considered to be linked to the experience of AVH. From Figures 2-4 and Table 2, it is quite obvious that the connections between the left and right superior parietal lobules as well as the left and right thalami seem to be substantially stronger during the HA condition. Attending to hallucinations also seem to reduce the strength of connections in the visual brain regions as well as the fronto-parietal network (Table 2). Another finding that was unique to the AVH+ sub-group was the reduction in connection strength between the bilateral superior temporal gyri.

There is a large body of work that discusses hallucinations as a deficit in action-monitoring or self-agency (Blakemore et al., 2002; Blakemore and Frith, 2003; Feinberg, 1978; Feinberg and Guazzelli, 1999; Frith, 1987; Frith and Done, 1988). Self-agency is the sense of authorship of one’s actions (Kircher and Leube, 2003). Detection of whether sensory signals are the result of self-generated actions or that of external agencies has been demonstrated to be effected by prediction of the sensory consequences of the self-generated motor act (Sperry, 1950; Wolpert et al., 1995; Wolpert and Flanagan, 2001) through a mechanism variously referred to as ‘efference copy mechanism’ (von Holst, 1954) or ‘corollary discharge mechanism’ (Frith, 1992; Sperry, 1950). Frith (Frith, 1992; Frith and Done, 1988) proposed the existence of a “cognitive self-monitoring” circuit based on the proposal by (Feinberg, 1978) that psychotic thinking could be linked to an impairment of the corollary discharge mechanism accompanying conscious thought. Impairment of the above corollary discharge mechanism accompanying conscious thought secondary to aberrations in the above circuit has been proposed to underlie symptoms such as hallucinations and delusions of alien control (Fu and McGuire, 2003).

The parietal lobe is the implementation site of the body-image and body-centred spatial concepts (Berlucchi and Aglioti, 1997; Melzack et al., 1997; Vogeley, 2003). Previous studies have shown that activity in the parietal cortex is typical of passive movements as opposed to active ones (Weiller et al., 1996). Moreover, the superior parietal lobe has been shown to be crucially involved in the awareness of self-agency (MacDonald and Paus, 2003). Spence et al., (1997) have shown over activity of the superior parietal cortex in patients with schizophrenia having delusions of alien control. Thus, the capacity to distinguish between self and others in these individuals may be deranged by a possible increased cortical ‘associativity’ (‘hyperassociativity’) (Behrendt, 1998) between the bilateral superior parietal lobules, apart from the prefrontal-parietal connections, which might lead to an imbalance between self-monitoring and reality modelling domains (Vogeley, 2003). This, in turn, results in an inability to recognize the origin of thought as being from oneself and a subsequent misattribution to external sources (Frith, 1992). Our finding of stronger connectivity between the bilateral superior parietal lobules that is linked to the experience of auditory hallucinations provides support to the above hypothesis of the neural substrate of auditory hallucinations. The left superior parietal lobule showed relatively stronger connection with the left inferior parietal lobule in the overall schizophrenia sample as well as the AVH+ subgroup during the HA condition (Table 2). Inferior parietal lobule mediates bottom-up reflexive reorienting of attention to behaviourally relevant information (Cabeza et al., 2008) Thus, the stronger connection between these regions might reflect the greater attentional allocation to auditory hallucinations during the HA condition as opposed to lesser attentional allocation given to the visual stimuli (see below).

Thalamus is the major relay center of the brain between different subcortical areas and the cerebral cortex. The medial geniculate nucleus of the thalamus acts as a key auditory relay center between the inferior colliculus of the midbrain and the primary auditory cortex. This allows the thalamus to play the role of an information ‘filter’ or sensory ‘gate’ (Andreasen, 1997). Many previous studies have reported decreased thalamic activations in patients with schizophrenia during rest or during performance of certain cognitive tasks (Andreasen et al., 1996, 1994; Buchsbaum et al., 1996). However, these studies were not designed to examine the brain activations associated with the experience of auditory hallucinations. Silbersweig et al., (1995) reported abnormal activations in thalamus in patients with schizophrenia during active auditory hallucinations. Our finding of increased functional connectivity between the bilateral thalami might indicate an increased effort by the thalamus to ‘filter’ out the aberrant auditory perceptual experiences in patients with schizophrenia that may be caused by top-down processing abnormalities involving the superior parietal lobule and its fronto-parietal connections.

The bilateral precuneus showed interesting connectivity patterns during the three conditions in healthy, schizophrenia and AVH+ subjects. Precuneus, along with the posterior cingulate gyrus is suggested to be the core hub of the default mode network that is active during resting state, thereby suggesting that it is an important brain region underlying conscious information processing (Cavanna, 2007; Vogt and Laureys, 2005). Wallentin et al., (2006) have reported that precuneus is involved in memory tasks requiring subjects to respond to questions regarding spatial details of images that were presented. In the present study, functional connectivity between bilateral precuneus was increased in the VA condition compared to the FA condition in both healthy *[Z values (FA/VA) =1.1267 /1.2346]* and schizophrenia *[Z values (FA/VA) =1.0729 /1.2676]*. Most notably, the AVH+ sub-group did not show significant functional connectivity between bilateral precuneus in the HA condition. This might reflect a shift of focus away from processing of the visual stimuli (associated with increased connectivity of the medial parietal lobe, the precuneus) to the experience of AVH (associated with increased connectivity of the lateral parietal lobe, the superior parietal lobule and a decreased connectivity of the bilateral precuneus), specifically in patients who experienced auditory hallucinations during the HA condition. This shift of focus away from the visual stimuli to the experience of auditory hallucinations might also explain the lack of significant bilateral insular as well as right anterior cingulate-medial prefrontal cortical connectivity as well as weaker bilateral anterior cingulate connectivity in the AVH+ subgroup, since, the visual stimuli may not be considered as salient during the HA condition.

Reduced fronto-temporal connectivity has been consistently reported in schizophrenia, especially in those with auditory hallucinations (Ford and Mathalon, 2005; Lawrie et al., 2002). In the present study, the connection strength between brain regions of the frontoparietal network was found to be substantially weaker during the hallucination attention condition as compared to the other conditions in both groups of subjects. In the AVH+ subgroup, the frontal connections were non-significant. This could indicate lesser attention allocated towards the visual stimuli while attending to hallucinations and a greater attentional allocation given to the hallucinations, as reflected by stronger left SuPL-left InPL connection strengths (see above). The same explanation could hold true for the observation of nonsignificant connections between the visual brain regions during the HA condition in AVH+ subjects.

The superior temporal gyrus (SuTG), which is the primary auditory area of the brain, has been implicated in the experience of AVH in several studies (for example, (Allen et al., 2007; Dierks et al., 1999; Sommer et al., 2008)). Significant connections were noted between the bilateral superior temporal gyri during FA and VA conditions in both groups of subjects, perhaps due to the fact that the connectivity between the SuTG homologues may have been uniformly increased in both groups due to the scanner noise. Interestingly however, the bilateral SuTG connections were substantially weaker during the HA condition in the AVH+ sub-group, perhaps because the attention was focussed in these subjects away from the scanner noise to the experience of the auditory hallucinations. This is in keeping with the results of Gavrilescu et al., (2010) who found reduced interhemispheric connectivity between the primary and secondary auditory cortices in patients with hallucinations, as compared to patients without hallucinations and healthy subjects.

Significant functional connections between bilateral medial prefrontal cortical as well as bilateral parahippocampal cortices were observed during all the three conditions in both groups of subjects, indicating their relevance for processing of the visual stimuli. However, there were no alterations of these connections in patients with schizophrenia or in the AVH+ sub-group, indicating that these connections may not be directly/indirectly linked to the experience of auditory hallucinations.

Thus, using functional connectivity analysis of fMRI time series acquired during the performance of a novel Hallucination Attentional Modulation Task (HAMT), we demonstrate the brain substrates of normal visual perception/processing in healthy and schizophrenia subjects; and the connectivity aberrations that are associated with the experience of AVH in patients with schizophrenia (Figure 5).

**Figure 5:**
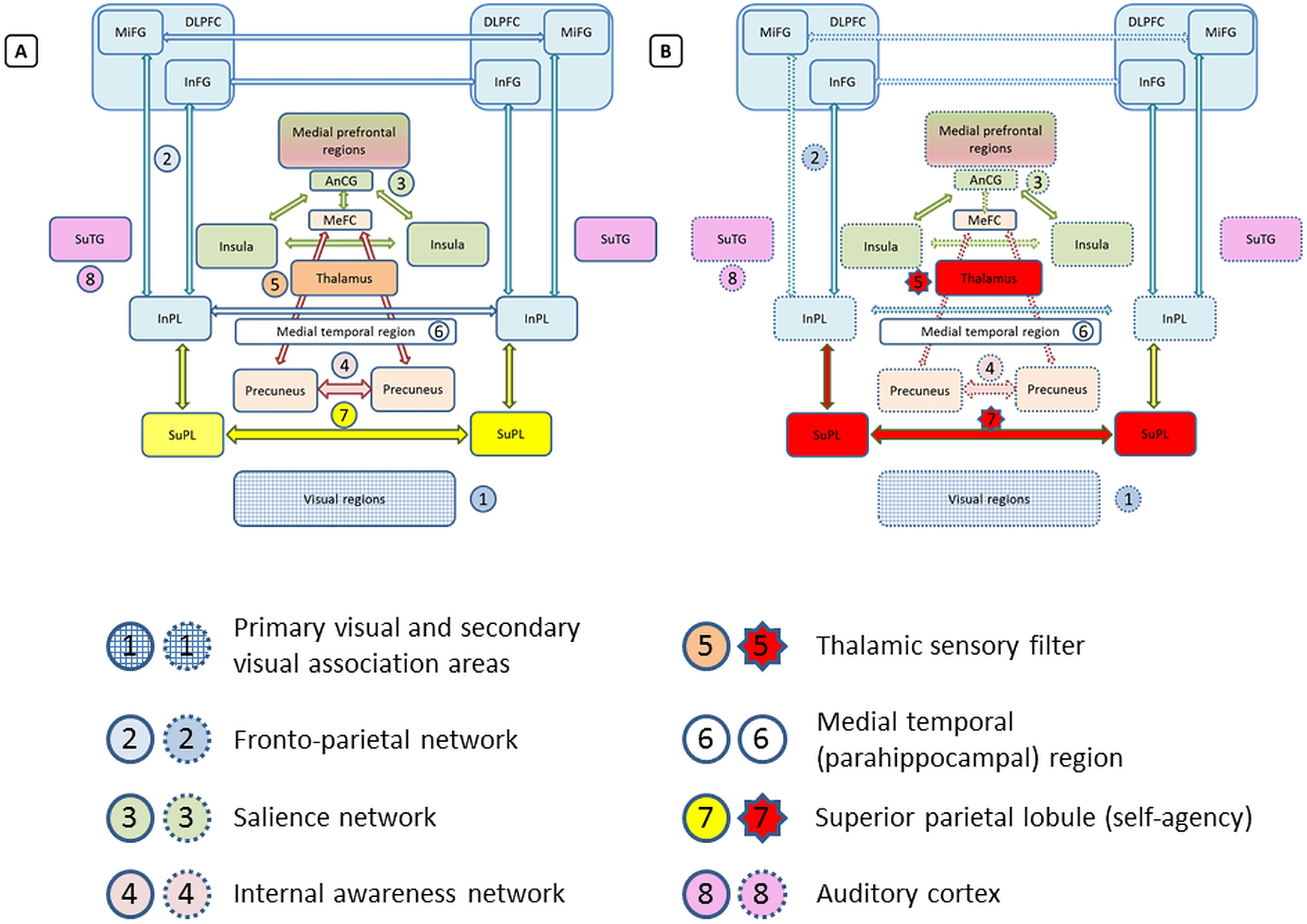
A schematic representation of the different brain networks/regions involved in the processing of ‘real’ visual stimuli in healthy and schizophrenia subjects (A) and the experience of auditory verbal hallucinations in patients with schizophrenia (B). The brain regions shown in the same colour connected by arrows in the same colour form part of the same network (refer the labels against the numbers given below the figure) that are involved in visual processing and auditory hallucination. In (B), brain regions/connections that showed increased connectivity during the experience of auditory verbal hallucinations are shown in red, while those regions/connections that showed weaker connections during verbal hallucinations are represented using dotted lines. *Note:* The superior temporal gyri (8) shown in (A), even though not an integral part of the brain regions involved in visual processing, showed strong connections during free and visual attention in both healthy and schizophrenia subjects probably as a result of the background scanner noise; however, the connection strength between the bilateral superior temporal gyri was noted to be weaker (shown in B) in the subgroup of patients who experienced auditory hallucinations during the ‘hallucination attention’ condition of the Hallucination Attentional Modulation Task (AVH+).

### Strengths and limitations

The major strengths of the study include the stringent criteria used for recruiting patients with continuous AVH and the objective method of identifying patients who experienced hallucinations during the fMRI acquisition. The HAMT paradigm that we have developed has several advantages over other methods that have been used in fMRI studies so far. The fMRI paradigm was designed in such a way that the fMRI time series during each task including the hallucination attention task were of equal length. Previous task designs that involved button press to indicate onset of hallucination would result in unequal lengths of the time series which can influence the results of connectivity studies. Functional connections in the schizophrenia sample that are most prominent during the HA condition, which become less prominent during the FA condition and get further attenuated or become non-significant during the VA condition may be considered as state-related aberrant activations associated with auditory hallucinations in schizophrenia. Moreover, the proposed task permits direct comparison between the brain activations/deactivations associated with hallucinations and those associated with a real percept in an alternate sensory modality (visual, in contrast to auditory). However, given the limited sample sizes in this pilot study, no definitive inferences can be made from the observations of this study, especially since we have not attempted to perform between-group comparisons of functional connectivity in schizophrenia and healthy groups during the different conditions. Thus, the results are to be considered preliminary at best, and a more robust study with larger sample sizes is required for confirmation of the findings.

## 5 CONCLUSIONS

The present study provides preliminary evidence in support of the utility of a novel fMRI paradigm, viz., Hallucination Attentional Modulation Task (HAMT) to detect the functional connectivity patterns underlying normal visual perception as well as the aberrations in functional connectivity during the experience of AVH in patients with schizophrenia. The functional connectivity patterns associated with processing of visual stimuli point towards involvement of brain regions that constitute the internal awareness network, external awareness/fronto-parietal network, salience network as well as medial temporal regions, visual brain regions, thalamus and posterior parietal regions. The experience of AVH was found to be associated with aberrations in the above connectivity patterns that involve substantially stronger connections between bilateral superior parietal lobules and bilateral thalami, weaker connections in the fronto-parietal, visual brain regions and bilateral superior temporal gyri, along with a lack of significant connections between the bilateral precuneus. The findings of our study point towards the possibility of impaired self-agency underlying the experience of AVH, and provides preliminary evidence for superior parietal lobule being the neural substrate of the same.

## 6 ACKNOWLEDGEMENTS

This work was supported by the Department of Biotechnology, Government of India (to JPJ; BT/PR/8863/MED/14/1252) and the National Institute of Advanced Studies (NIAS) - Mani Bhaumik Research Fellowship (to PP; NIAS/MBRF-07/2014). We would like to acknowledge Dr. Alfonso Nieto-Castanon for his inputs at various stages of analyses. Part of this work was presented as a poster at the International conference on Cognition, Culture, and Consciousness at NIAS (9-11^th^ December, 2015) and also appeared as an abstract of the proceedings of the Brain Informatics and Health conference (30^th^ August – 2^nd^ September, 2015, London).

## 7 DATA AVAILABILITY

Data currently not available publically. However, the authors will be happy to share unthresholded statistical maps, atlases, etc. on public repositories like NeuroVault upon acceptance of the manuscript.

